# A Modular Microfluidic System to Generate Gradient Hydrogels with Simple-to-Complex Stiffness Profiles for Mechanobiology

**DOI:** 10.1101/2025.05.23.655778

**Authors:** Shin Wei Chong, Jameel Sardharwalla, Gweneth Sofia P. Masonsong, Joshua Cosgrove, Anthony Katselas, Isaac J. Gresham, Marcela M.M. Bilek, Yi Shen, Chiara Neto, Daniele Vigolo

**Affiliations:** School of Biomedical Engineering, The University of Sydney, Sydney, NSW 2006, Australia; School of Chemistry, The University of Sydney, Sydney, NSW 2006, Australia; School of Physics, The University of Sydney, NSW 2006 Australia; School of Chemical and Biomolecular Engineering, The University of Sydney, Sydney, NSW, 2006, Australia

**Keywords:** stiffness gradient hydrogels, complex gradient, microfluidics, thermophoresis, mechanobiology

## Abstract

Engineered stiffness gradient hydrogels offer exciting opportunities to probe fundamental mechanobiological processes *in vitro*. However, the need to spatially manipulate the properties of soft hydrogels at the micron scale poses challenges in developing fabrication platforms that can reliably modulate the gradient gel characteristics according to user needs. This study describes a modular approach leveraging thermophoresis in microfluidics to create high-fidelity stiffness gradients with linear and complex profiles, including periodic and anisotropic gradients. This study describes the platform’s design and optimization, demonstrating achievable stiffness ranges and gradient slopes that correlate with many physiologic and diseased tissue types. This platform is also compatible with different hydrogel crosslinking chemistries, providing a versatile tool to engineer microenvironments with increased complexity. Directionally biased fibroblast cell proliferation and migration on fabricated stiffness gradient gelatin methacryloyl (GelMA) hydrogels indicate the effectiveness of this platform in modulating the mechanical microenvironment of cells. The results indicate that both the absolute stiffness range and the pattern of stiffness variation jointly affect cell behaviors. Considering its remarkable flexibility, the fabrication platform can significantly advance the development of biophysical gradient hydrogels that better replicate the intricacies of native tissues and help realize the next breakthroughs in mechanobiology.

## 1. Introduction

Microscale physical gradients are desirable features in hydrogels to increase their biological relevance for mechanobiology and tissue engineering applications, prompting substantial research efforts. Given that the extracellular matrix (ECM) stiffness is known to be a major regulator of cell behavior,^[1,2]^ numerous advances have been made in stiffness gradient hydrogels to mimic the heterogenous tissue mechanics *in vivo*,^[3–5]^ or as a screening tool to probe cell-material interactions on a continuum of conditions.^[6–8]^ While a range of hydrogel systems have been designed to replicate the native ECM *in vitro*,^[9,10]^ stiffness gradient hydrogel platforms are intentionally simple to decouple the complexity of the cellular microenvironment. These platforms can be a powerful tool in driving our mechanistic understanding of how cells sense the physical cues in their surrounding matrix, with implications in cell signaling and how to modulate cell behavior by programming the ECM.

The basic premise for biomimicry of stiffness gradient hydrogels is that they represent the stiffness range and gradient strength characteristic of the target tissue type or biological process. In this regard, the stiffness (defined in this study as the **Young’s modulus)** of tissues spans orders of magnitude, from a few kilopascals (kPa) for soft tissues such as the brain to hundreds of kPa for osteochondral tissues.^[11]^ On the other hand, the required stiffness gradient strength can range from ≈1 kPa mm^−1^ in physiological tissues to ≈10 kPa mm^−1^ in pathological tissues and >100 kPa mm^−1^ at tissue interfaces.^[12]^ To date, various strategies have been developed to create microscale stiffness gradients *in vitro*, including the commonly employed patterned photopolymerization,^[6,13,14]^ microfluidic gradient generators,^[15,16]^ and two-step polymerization techniques.^[5,17]^ Yet, each technique has its own tradeoff between achievable stiffness ranges, generated gradient shape fidelity, and ease of operation.^[18]^ Consequently, the goal of achieving a universal platform capable of interrogating the full physiological mechanical landscape while maintaining precise control of the gradient profile at single-cell length scale has remained an open challenge (**Table S1**).

Although conceptually straightforward, there are significant technical challenges in fabricating precise and reproducible gradients across biologically relevant stiffness ranges. Most current prevalent techniques are based on passive molecular diffusion,^[4,17,19]^ which imposes limited control over the gradient shape fidelity and resolution due to its sensitive dependence on the timing of each process step and external environmental factors such as temperature and humidity. The other main class of gradient generation techniques involves varying the crosslinking degree, typically using graded exposure to heat or light.^[13,14,20,21]^ Despite the possibility of achieving well-defined, steep, or complex gradients, these techniques generally require specialized equipment and careful management of process deviations introduced by equipment variability to ensure consistency.

Motivated by the vision of a gradient hydrogel platform that can be streamlined for a wide range of hydrogels and biological applications, we pioneered a fundamentally new approach that leverages the thermophoretic drift of hydrogel precursor molecules along a temperature gradient to generate concentration gradients within an initially homogenous solution.^[22]^ Since thermophoresis works by manipulating the precursor distribution in the liquid phase, this technique can be universally applied to nearly all hydrogel types independent of the crosslinking chemistry. We previously showed the implementation of thermal microfluidic devices to characterize microscale thermophoresis^[23]^ and their potential utility to locally tune the physical properties in hydrogels,^[22,24]^ while demonstrating their compatibility for *in vitro* cell culture. However, in our earlier studies, the microfluidic systems were designed to work exclusively with specific hydrogel types, limited to thin (up to ≈600 µm in width) strips of linear gradient gels, and involve a cumbersome setup and sample retrieval step, limiting widespread adoption.

To address these issues, here we introduce a modular microfluidic system that is universally applicable for producing linear or complex stiffness gradients (**Figure 1**). The platform is based on thermophoresis to achieve controlled generation of microscale gradients. In contrast to previous studies, however, this platform employs a two-part design that separates hydrogel processing from the microfluidic structure. Using this strategy, the microfluidic device is used to control local temperature differences within the platform – a requisite for thermophoresis process. The hydrogel sample would be contained in an external microchamber interface with the microdevice, which overcomes key challenges such as enabling easy hydrogel handling, reducing fabrication turnover time, and facilitating reusability of the thermal patterning microdevices. In this respect, the modular fabrication platform approaches the simplicity of mold casting methods.^[17]^ Furthermore, the modularity of the microfluidic thermal patterning module allows users to customize the stiffness gradient hydrogel features based on their experimental requirements, offering superior flexibility compared to current conventional methods.

**Figure 1:**
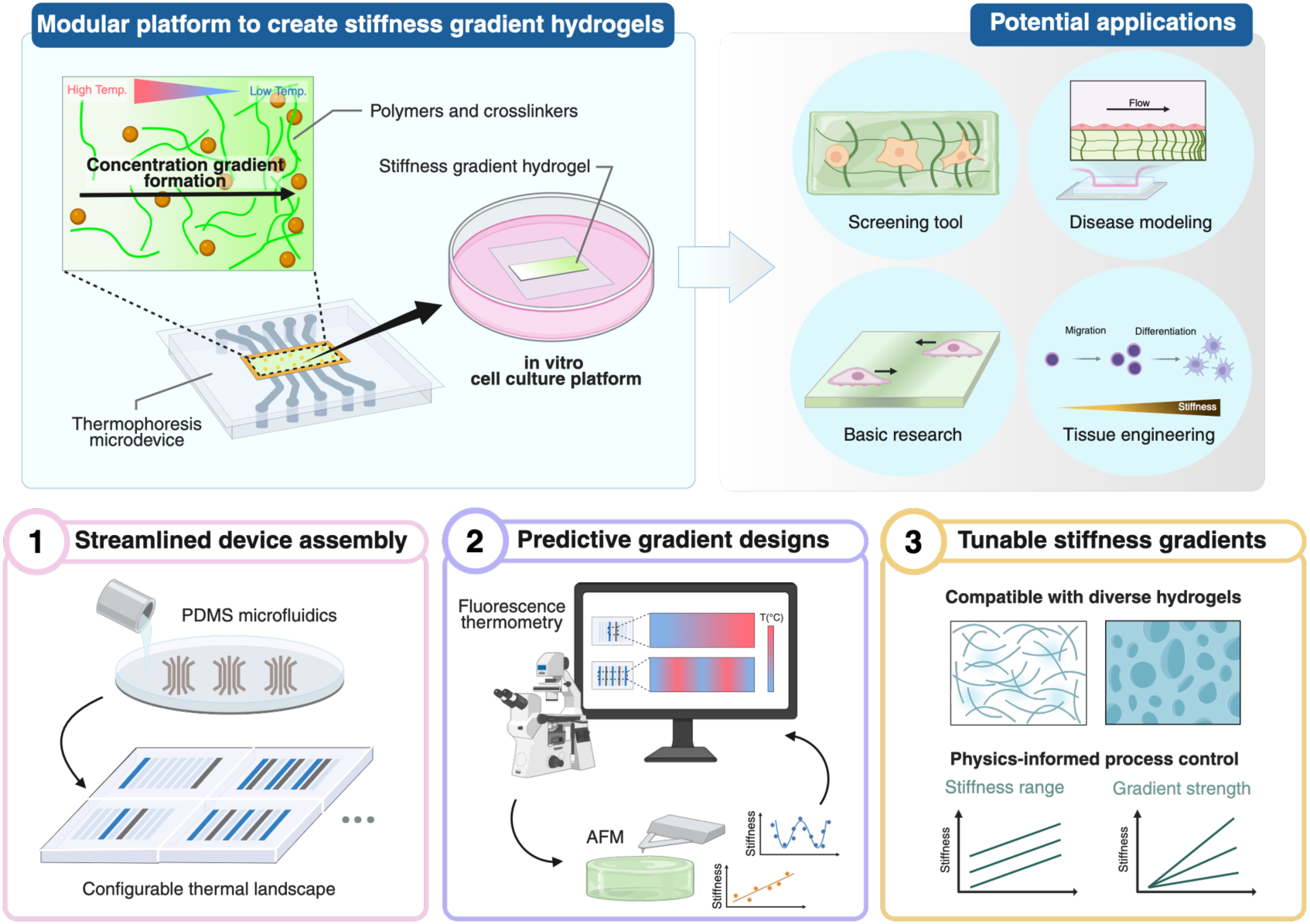
Conceptual illustration of a modular platform utilizing thermophoresis to fabricate stiffness gradient hydrogels for *in vitro* cell culture applications. (1) This approach leverages simple PDMS casting to rapidly produce microfluidic thermal patterning modules with customizable heating configurations. These configurations define the microscale temperature landscape within the system, which directly governs the resulting stiffness gradient profile in the hydrogel. (2) The generated temperature profile can be experimentally mapped at high spatial resolution using a fluorescent-based temperature probe. This information provides a predictive framework for users to optimize the module design and process parameters, enabling precise control over the stiffness gradient characteristics. (3) The flexibility, predictability and rapid design-to-fabrication cycle of the modular gradient hydrogel platform offers a scalable solution to study cell-material interactions across a wide range biologically relevant mechanical microenvironments, with promising future applications in biomaterials-based cell studies and tissue engineering.

This study shows the generation of linear, periodic, and anisotropic (sawtooth) gradients, demonstrated using thermosensitive Gellan gum (GG) and photopolymerizable gelatin methacryloyl (GelMA). Through extensive experimental characterization, it was established that the modular microfluidic device offers remarkable temperature control down to microscale resolution, underscoring its functionality to create reproducible stiffness gradient hydrogels with exceptional fidelity. The stiffness range and gradient strength are tunable by varying the initial polymer concentration and the strength of the applied temperature gradient, respectively, a feature of the thermophoretic process found to be general across different hydrogel types. The fabrication of GelMA hydrogels with various gradient profiles was achieved, and mouse fibroblast (3T3-L1) cells adjusted their proliferation and migration behaviors according to the different stiffness patterns. Overall, this work establishes a versatile approach for microscale patterning of soft functional biomaterials, which could potentially help realize the next advancements in mechanobiology and enable new applications in tissue engineering.

## 2. Results

### 2.1. Overview of the Gradient Hydrogel Fabrication Platform

The gradient hydrogel fabrication platform introduced in this work consists of a microfluidic thermal patterning module and a detachable hydrogel sample microchamber (**Figure 2**). The thermal patterning module is fabricated using polydimethylsiloxane (PDMS) and comprises nine equally sized microchannels in parallel (**Figure S1**). Each channel can be selectively opened and employed for cooling or heating, enabling a high degree of user-defined flexibility in configuring the microscale thermal landscape. To build a complete assembly, the PDMS module is plasma-bonded to a glass coverslip, on top of which a microchamber made by stacking rectangular cutouts of polymide (Kapton) adhesive tape is attached. Given the open nature of the microchamber, hydrogels can be fabricated using this platform in a similar manner to now-standard spacer techniques,^[14]^ greatly simplifying its operation.

**Figure 2:**
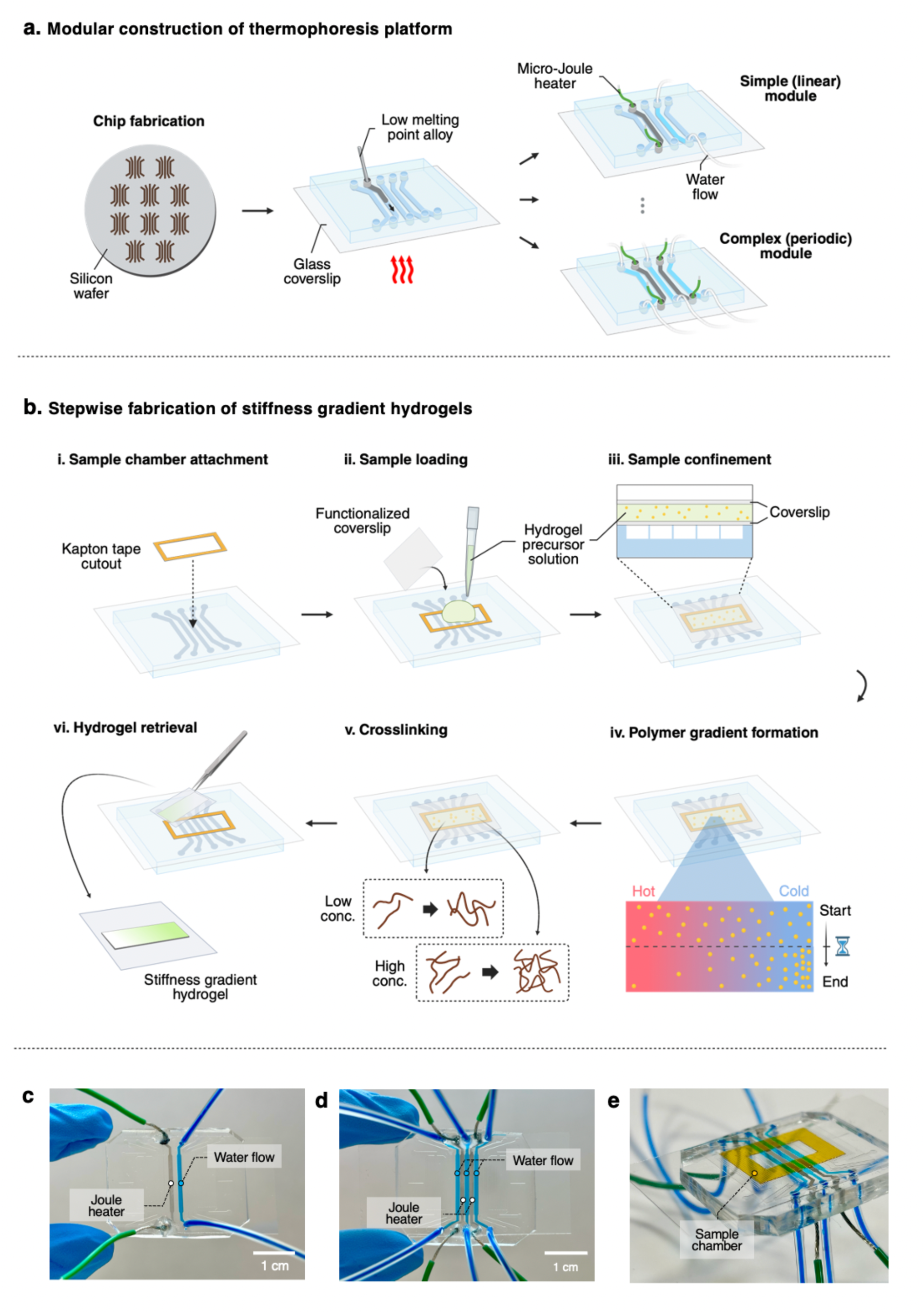
(a) Schematic overview of the thermophoresis microdevice fabrication via PDMS soft lithography, followed by formation of micro-Joule heaters by injecting low melting point alloy into the microchannels. Custom gradient patterning modules are made by configuring the arrangement of heating (micro-Joule heater) and cooling (water flow) channels. (b) Stepwise demonstration of stiffness gradient hydrogel fabrication using the thermophoresis platform. (i) A Kapton tape cutout is attached to define the sample chamber. (ii) Hydrogel precursor solution is added into the chamber, and (iii) a chemically treated coverslip is placed on top to enable hydrogel attachment. (iv) Under an applied temperature gradient, microscale concentration gradient of precursor molecules in the solution forms via thermophoresis effect. (v) After a specified duration, the polymer solution is rapidly crosslinked and (vi) removed from the microdevice to result in a stiffness gradient hydrogel. Representative images of the thermophoretic fabrication platform, including (c) a simple linear gradient module, (d) a complex periodic module, and (e) the complete platform assembly with a sample chamber.

The modular microfluidic system operates under the same physical principles of thermophoretically driven gradient generation as described previously.^[22,24]^ Cooling is achieved using a syringe pump to drive fluid through the microchannel, whereas heating is achieved by embedding a micro-Joule heater, operated by DC currents, in the corresponding channels. Specifically, the micro-Joule heaters are created using low melting point alloy (LMPA) that flows into the microchannel via capillary action when heated above its melting point (≈98 °C), forming a conductive path once it cooled and solidified (**Figure 2a**). When the hydrogel precursor solution, confined in the microchamber, is subjected to spatially patterned temperature from the underlying microfluidic module, the solutes accumulate toward either the cold or hot regions of the system, depending on the Soret coefficient (*S_T_*) (**Figure 2b**).^[22,25]^ For the hydrogel systems explored in this study, accumulation occurred from the hot to the cold regions, indicating a positive *S_T_*. The degree of accumulation varies depending on the magnitude of the applied temperature gradient and the average operational temperature, which in turn determines the local value of stiffness upon hydrogel crosslinking. The stiffness of the hydrogels, characterized by the Young’s modulus, generally increases proportionally with the polymer or crosslinker concentration.^[26]^ As a result, for the systems considered in this study, the generated stiffness gradient profile is an inverted replica of the predefined thermal pattern. Using this thermophoretic technique, the initial polymer concentration and temperature gradient strength were tuned to achieve the desired outcomes of a stiffness gradient gel. Throughout this work, the devices were operated at a minimum temperature gradient of 7 °C mm^−1^ to rapidly and effectively generate a precursor concentration gradient. An average system temperature of above 60 °C was also used to reduce solution viscosity and prevent premature gelation of GG and GelMA. Importantly, thermophoresis occurs whenever a temperature gradient is maintained, irrespective of the specific temperature range.^[25]^ This property makes the platform universally applicable to gradient gel fabrication using other hydrogel types that have relatively low viscosity at milder temperatures, including polyacrylamide and collagen, which is desirable for 3D cell encapsulation applications.^[27]^

Microfluidic modules with different stiffness patterning capacities were designed to suit the needs of various applications. In the simplest case, linear gradient devices are built with a single pair of cooling and heating channels (**Figure 2c**). The thermophoresis characteristic time is consistent with the diffusion characteristic time following the Stokes-Einstein relation.^[24]^ Thus, one could expect that the steepest gradients are created by utilizing adjacent pairs of channels in the microchannel array, while shallower gradients are created with increasing separation between the channels. Since it is possible to independently generate multiple different linear temperature profiles that blend at the transition points, various complex gradients can be established over ≈10 mm in the microfluidic system (**Figure 2d**). Under essentially the same thermophoresis principles, solutes in the precursor solution distribute in the direction of the temperature gradient and proportionally to the gradient strength, resulting in a smooth stiffness gradient profile. Although existing microfluidic gradient generators can create similarly well-defined microscale stiffness gradients, these devices are typically single-use only and a unique microfluidic device design is required for a specific gradient shape and size, which makes this technique costly and time-consuming.^[28]^ These issues can be solved using the modular approach presented here, because the assembled microdevices can be reused multiple times, requiring only replacement of the microchamber spacer between uses (**Figure 2e**).

### 2.2. Temperature Characterization of The Microfluidic System for Predicting Gradient Hydrogel Properties

The fabrication of precise and reproducible stiffness gradients using the modular thermophoretic platform relies on the ability to generate and maintain intricate temperature patterns within the microsystem. A crucial innovation in this study is the implementation of a fluorescent-based temperature measurement method, specifically designed to enable high spatiotemporal resolution mapping of the temperature distribution in the microfluidic system (**Figure 3a**). This strategy employs a glass microcapillary filled with Rhodamine B solution as a temperature probe and leverages the temperature-dependent fluorescence of Rhodamine B^[29]^ to evaluate the local temperature based on changes in fluorescence intensity (**Figure S2**). The microcapillary was attached to the microfluidic module following a similar setup for microchamber attachment for hydrogel fabrication (**Figure 3b**). Thus, this setup effectively captures the temperature profile inside the microchamber under fabrication-relevant conditions, offering a potential powerful tool to assess the experimental design space and extract predictive process insights.

**Figure 3:**
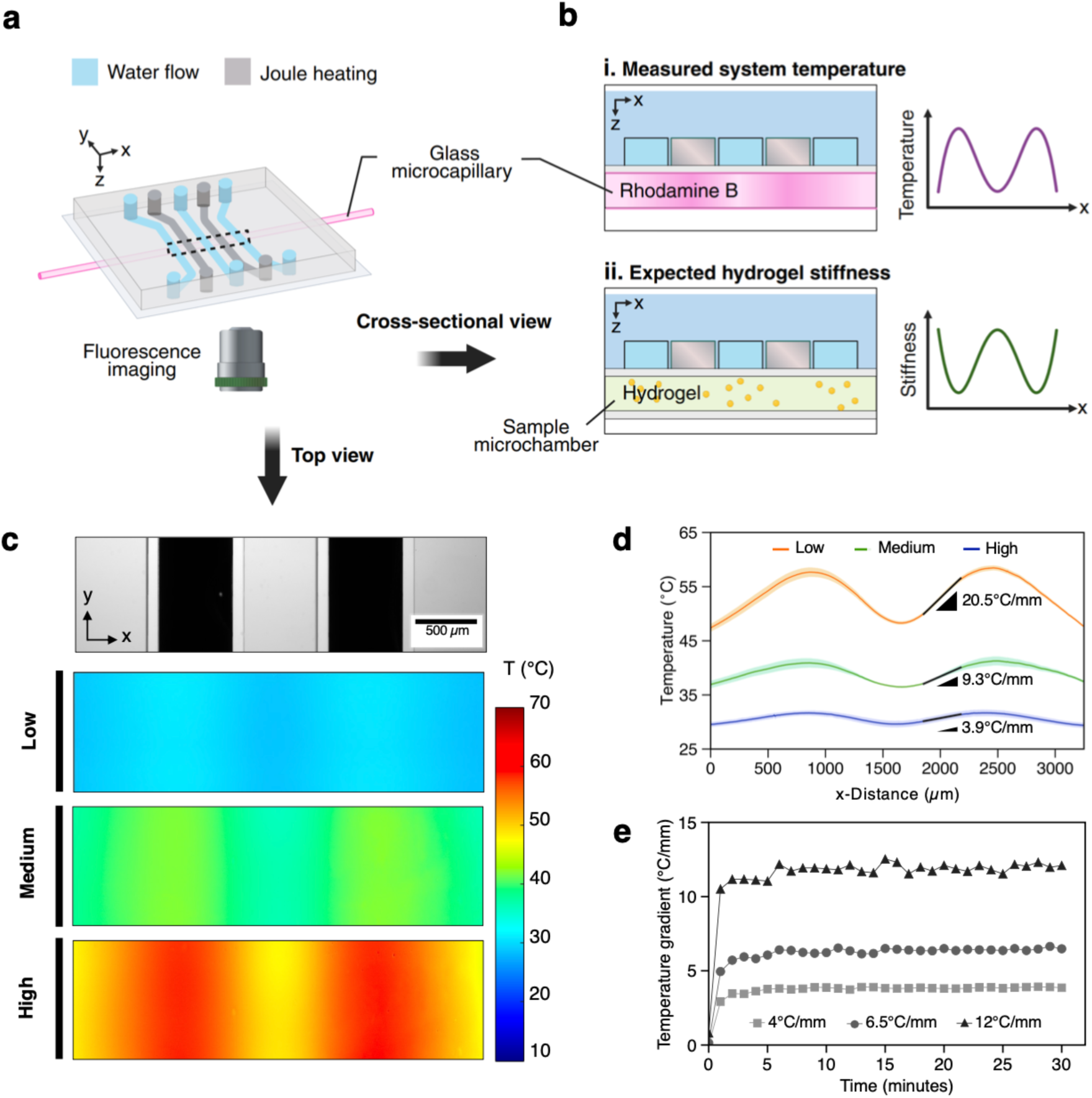
(a) Schematic of experimental setup utilizing temperature sensitive Rhodamine B dye to characterize temperature distribution within the microfluidic system. (b) Graphical illustration of the system cross-section comparing the placement of (i) microcapillary for temperature characterization and (ii) microchamber for gradient hydrogel fabrication. The Rhodamine B-filled microcapillary enables fluorescent-based measurement of the temperature profile representative of the conditions within the microchamber, and precisely mirrors the resulting hydrogel stiffness profile. (c) Optical micrograph showing the top view of a thermal patterning module configured with alternating water flow channel (bright lane) and micro-Joule heater (dark lane), demonstrating wide-range temperature control with microscale precision. (d) The corresponding temperature profile of (c), obtained by averaging in the channel direction (y-axis). (e) The system temperature was controlled by varying the water flow rates and Joule heater current, demonstrating rapid and precise control over an extended duration.

To assess the platform’s capabilities for creating complex gradients, initial experiments were performed with a focus on a 5-channel microfluidic module designed for periodic gradients (**Figure 3**). Temperature characterization results confirmed the successful generation of a uniform periodic gradient across a wide range of temperature values and clearly demonstrated tunable temperature gradient slope (≈4 °C mm^−1^ to ≈20 °C mm^−1^) while maintaining the gradient shape fidelity (**Figure 3c,d**). Specifically, temperature control was achieved by altering the water flow flows or current through the Joule heaters (**Table S2**). Real-time analysis at various temperature gradient strengths further revealed that, upon initiating temperature control, the system rapidly (<2 minutes) approached the target temperature profile and remained stable for at least 30 minutes (typical thermophoresis process time for gradient hydrogel fabrication) (**Figure 3d**). Such precise temperature control of the platform enables “on-demand” and predictable progression of the thermophoresis process, which is essential to ensure highly reproducible stiffness gradients. It is important to note that these system behaviors were also found to be consistent regardless of the thermal configurations (**Figure 4,5**), highlighting the versatility of the modular microfluidic system to pattern microscale gradients in a highly controlled manner.

**Figure 4:**
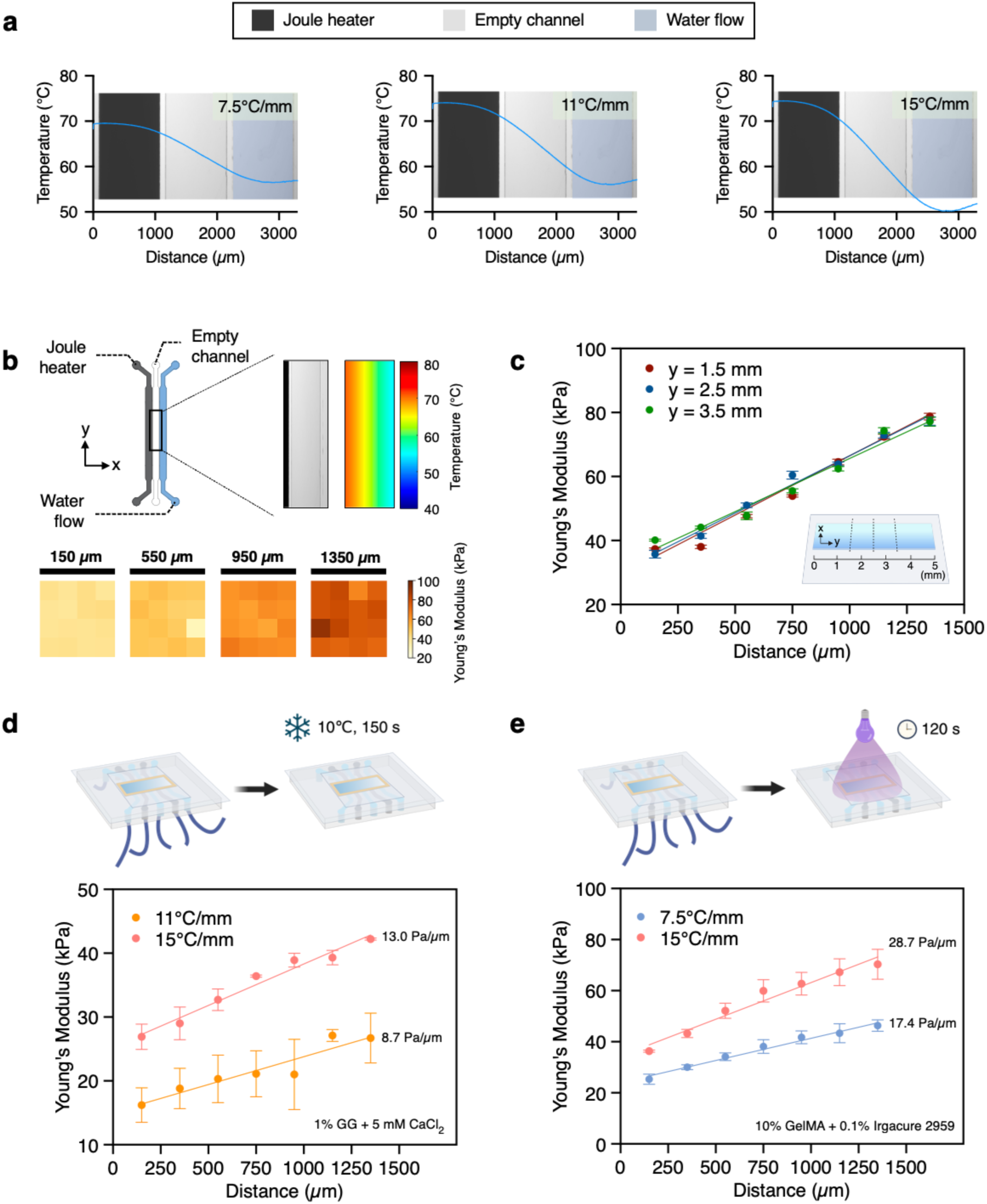
(a) Temperature measurement of the microfluidic system superimposed onto the brightfield image of the microdevice configured for generating linear gradients. (b) Representative platform setup for fabrication and characterization of linear stiffness gradient hydrogels. The black bounding box indicates the magnified view of the sample microchamber area measuring 1500 µm × 5000 µm (top). The micrograph shows the top view of the region of microdevice coinciding with the sample area, in the presence of a uniform temperature gradient of 15 ℃ mm^−1^ along the chamber length (top, right). Variation in stiffness was measured by taking AFM force maps (16-point grid, 20 µm × 20 µm area) at regular intervals across the gradient hydrogel (bottom). The numbers shown above each map indicate the distance from the hot end of the microchamber. (c) Stiffness measurements taken by AFM across the gradient hydrogel showed that stiffness varies gradually across the width of the sample (x-direction) and remains constant along the sample (y-direction). AFM measurement on (d) temperature-dependent crosslinked GG and (e) photopolymerized GelMA stiffness gradient hydrogels. Error bars represent standard error.

**Figure 5:**
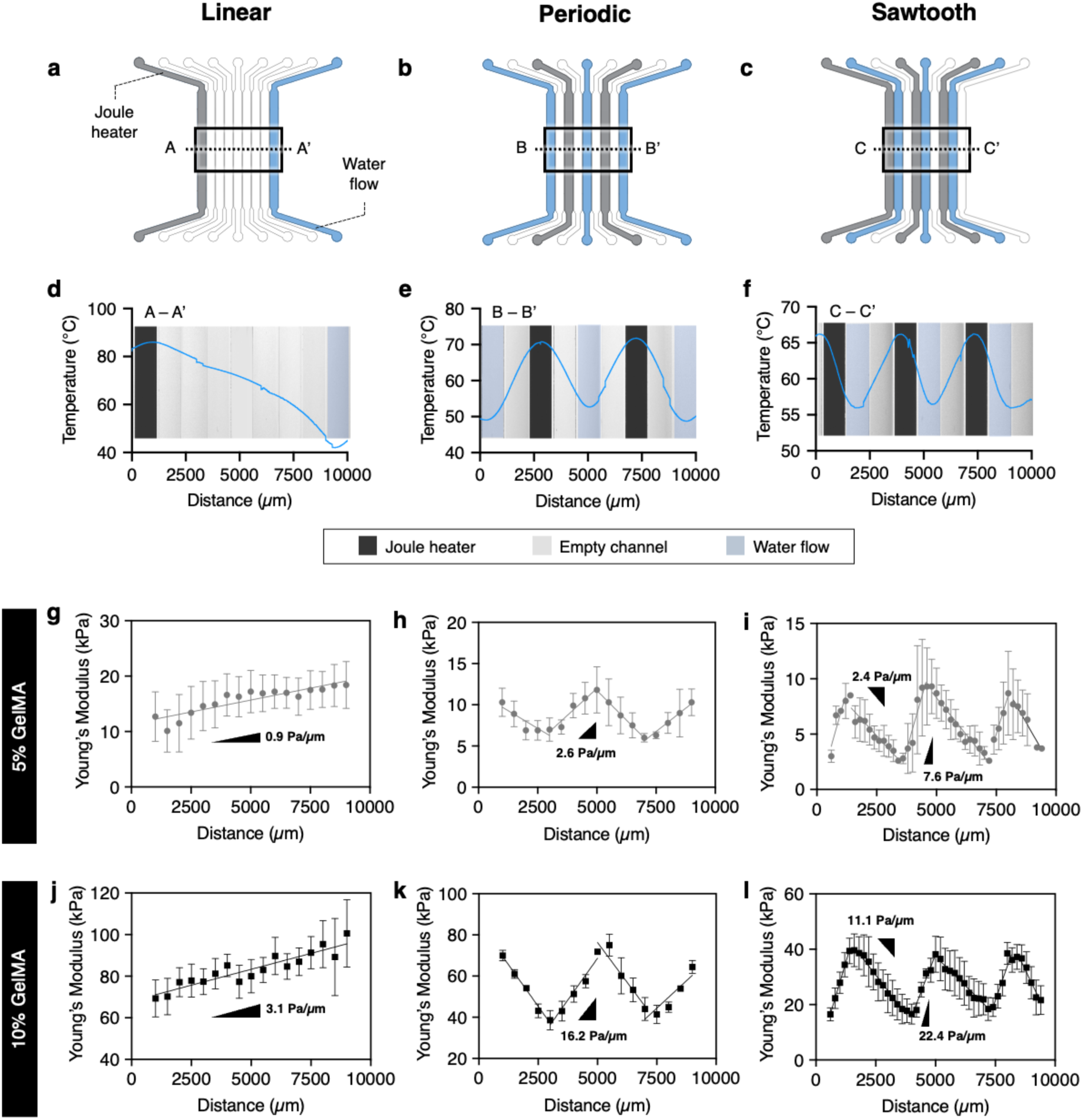
(a-c) Different gradient profiles can be achieved from a single PDMS microfluidic device by altering the configuration of heating (Joule heater) and cooling (water flow) channels. Black bounding boxes indicate the sample microchamber area measuring 10000 µm × 5000 µm. This approach of microfluidic thermal patterning creates a smooth temperature gradient profile, including (d) linear, (e) periodic, and (f) sawtooth shapes. AFM measurements of stiffness gradients in hydrogels fabricated using (g-i) 5% w/v GelMA and (j-l) 10% w/v GelMA show that the resulting stiffness profiles mirror the imposed temperature gradients. Error bars represent standard error.

Next, thermophoresis experiments using fluorescent polystyrene nanoparticles (fluo-NP) were performed to verify the underlying mechanism for creating gradient materials using the modular platform (**Figure S3**). Time-lapse imaging revealed the progressive formation of a continuous distribution of fluo-NP in response to the imposed temperature gradient, confirming the dominant effect of thermophoresis. Furthermore, to our knowledge, this study is the first demonstration of utilizing thermophoresis to generate spatially complex yet controlled concentration gradients within micro-confined geometries.

A key advantage of this modular microfluidic system design is that the measured temperature profile precisely reflects the resulting profile of thermophoresis-driven concentration gradients. In the context of hydrogel systems, the polymer concentration directly impacts the substrate stiffness, and thus, the imposed temperature profile can directly inform the expected hydrogel stiffness pattern (**Figure 3b**). Importantly, this unique feature allows users to use the fluorescent-based temperature characterization tool for high-throughput screening of different device configurations or process parameters to predict the final stiffness gradient characteristics. This contrasts to conventional workflows requiring iterative cycles of gradient gel fabrication and cumbersome AFM characterization to optimize the process conditions, enabling potential huge time savings. Future work could leverage the process input-output relationships to derive data-driven predictive models and design rules toward increasingly complex yet controlled gradient hydrogel systems.

### 2.3. Creating Linear Stiffness Gradients in Gellan Gum (GG) and Gelatin Methacryloyl (GelMA) Hydrogels

Next, the capability of the microfluidic platform to pattern microscale stiffness gradients in different hydrogel types was assessed. For these experiments, linear stiffness gradient hydrogels (1500 µm in width) were fabricated using three different temperature conditions: 7.5 °C mm^−1^, 11 °C mm^−1^ and 15 °C mm^−1^ (**Figure 4a**). To further demonstrate the compatibility of this platform with different hydrogel crosslinking schemes, GG and GelMA were selected for testing as representatives of thermally crosslinked and photopolymerized hydrogels, respectively. GG is a naturally occurring polysaccharide with attractive mechanical properties, making it a popular choice for osteochondral applications in tissue engineering.^[30]^ On the other hand, GelMA is a gelatin derivative that has been widely used for diverse biomedical applications due to the presence of desired cell-adhesion motifs, highly tunable biophysical properties, and versatility in terms of processing methods.^[26,31]^

Stiffness characterization using atomic force microscopy (AFM) was performed to evaluate the accuracy of stiffness patterning and the resulting stiffness gradient profiles. In the case of GelMA (10% w/v initial polymer concentration; 15 °C mm^−1^ applied temperature gradient), for example, the results show an expected linear stiffness gradient that closely matches the inverse profile of the temperature gradient (**Figure 4b**). AFM measurements at various length axes further revealed that the stiffness gradient was consistent across the whole gel (**Figure 4c**). These results suggest the effectiveness of thermophoresis for manipulating polymers in microvolumes and limited dispersion of the established concentration gradient during the crosslinking process. Importantly, the gradient gels exhibit a highly uniform local stiffness, as measured within a 20 µm x 20 µm area (AFM force maps in **Figure 4b**). This property is crucial to elucidate how cells respond to spatially organized mechanical signals on the micrometer to millimeter length scale, as opposed to variations at the subcellular level, which could have a confounding effect on cell-matrix mechanotransduction.^[32]^

Following the thermophoretic generation of a concentration gradient, rapid crosslinking is the next crucial step to preserve this gradient and ensure good reproducibility of the final stiffness values. GG crosslinking is achieved by cooling the microfluidic fabrication platform below the gelation temperature (**Figure 4d**), whereas GelMA is chemically crosslinked by exposing the entire platform to UV light (**Figure 4e**). From the AFM measurements, it is clear that GelMA hydrogels yielded a steeper stiffness gradient compared to GG hydrogels at the same applied temperature gradient (**Figure 4d,e**). Furthermore, at 7.5 °C mm^−1^, a measurable stiffness gradient was observed in GelMA but not GG hydrogels. It was found that an applied temperature gradient >10 °C mm^−1^ was necessary to generate a discernible variation in stiffness across GG hydrogels, which can be explained since dispersed GG fibers are significantly larger than GelMA monomers and thus require a stronger temperature gradient to induce the requisite thermophoretic force.^[33]^

Currently, it is common for researchers to employ several distinct fabrication methods to comply with the hydrogel choice or to cover the required stiffness landscape,^[12,34]^ each of which has its separate set of optimization criteria to create the desired results. Using the approach in this study, it is important to note that the optimal fabrication parameters might differ slightly between polymer systems due to inherent variations in their thermophoretic dynamics.^[32,35–37]^ However, the fact that the platform can be flexibly applied to different chemistries enables a modular approach for the development of stiffness gradient hydrogels which makes it more accessible to new users.

### 2.4. Tuning The Gradient Profiles via Microfluidic Thermal Patterning

While the aforementioned results focused on short-range, steep linear stiffness gradients, the study of mechanobiological processes requires cellular analysis in gradients of different stiffness ranges, gradient slopes, and lateral dimensions of the gradient (on the order of micrometers to centimeters).^[12,18]^ The microfluidic design was optimized to achieve a fine balance between the geometry and patterning resolution. To demonstrate the potential of the platform for creating a wider variety of gradient profiles, subsequent discussions will focus on GelMA hydrogels, which is justified given their growing popularity as cell culture substrates.^[15,38–40]^

By adjusting the configuration of heating and cooling channels within the microfluidic module, long-range, shallow linear gradients (< 5 Pa µm^−1^) spanning ≈10 mm in width can be generated (**Figure 5a,d,g,j**). Contrasting to the steep, pathological-mimicking gradients (>10 Pa µm^−1^) as shown in **Section 2.3**, these shallow gradients are useful to recreate certain tissue mechanical microenvironments, or in applications where durotactic behavior (i.e., cell migration towards stiffer or softer substrate regions) is undesired.^[6,17]^ Beyond monotonic gradients, the feasibility of generating complex stiffness patterns using the proposed platform was also demonstrated. Alternating heating-cooling channel configuration enabled periodic gradients (**Figure 5b,e,h,k**) as well as sawtooth gradients comprising a repeating period of steep positive slope and shallow negative slope (**Figure 5c,f,i,l**). In all cases, the resulting stiffness profiles were in good agreement with empirical predictions based on the temperature characterization results. Moreover, tuneability in stiffness values can be achieved by adjusting the initial polymer concentration without compromising the resulting gradient shape fidelity (**Figure 5g-l**; 5% w/v GelMA vs. 10% w/v GelMA). In this study, the heating and cooling channels were connected in parallel such that the system can be operated using a single power supply unit and syringe pump (**Table S2**). However, given that each microchannel can be controlled separately, it is possible to extend the accessible range of gradient profiles with added complexity. Future promising areas of application include advanced *in vitro* platforms for mechanobiology research, integration into organ-on-a-chip models for improved physiological relevance, and mechano-guided design of biomaterials for tissue engineering.^[41,42]^

### 2.5. Fibroblast Cell Behaviors on Varying Stiffness Gradient GelMA Hydrogels

Having established that the microfluidic manipulation of thermophoresis can be used to fabricate simple linear and complex stiffness gradient hydrogels, 3T3-L1 fibroblasts were seeded on these gradient hydrogels to assess the potential of this gradient hydrogel platform for studying stiffness-dependent cellular responses. To this end, 10% w/v GelMA system was selected as the base material to create three distinct stiffness profiles: a steep linear gradient (**Figure 6c**; stiffness ≈35-80 kPa, slope ≈28 Pa µm^−1^), a periodic gradient (**Figure 6e**; stiffness ≈40-80 kPa, slope of linear part ≈16 Pa µm^−1^), and a dual-slope gradient produced using a single period on the sawtooth pattern module (**Figure 6h**; stiffness ≈15-40 kPa, slope of shallow part ≈11 Pa µm^−1^, slope of steep part ≈22 Pa µm^−1^). In all cases, the hydrogel samples were used directly for cell culture experiments without additional ECM protein coating (**Figure 6a**). Cell nuclei and the actin cytoskeleton were stained for visualization, which were subsequently divided into multiple segments parallel to the direction of the gradient for quantitative cell analysis.

**Figure 6:**
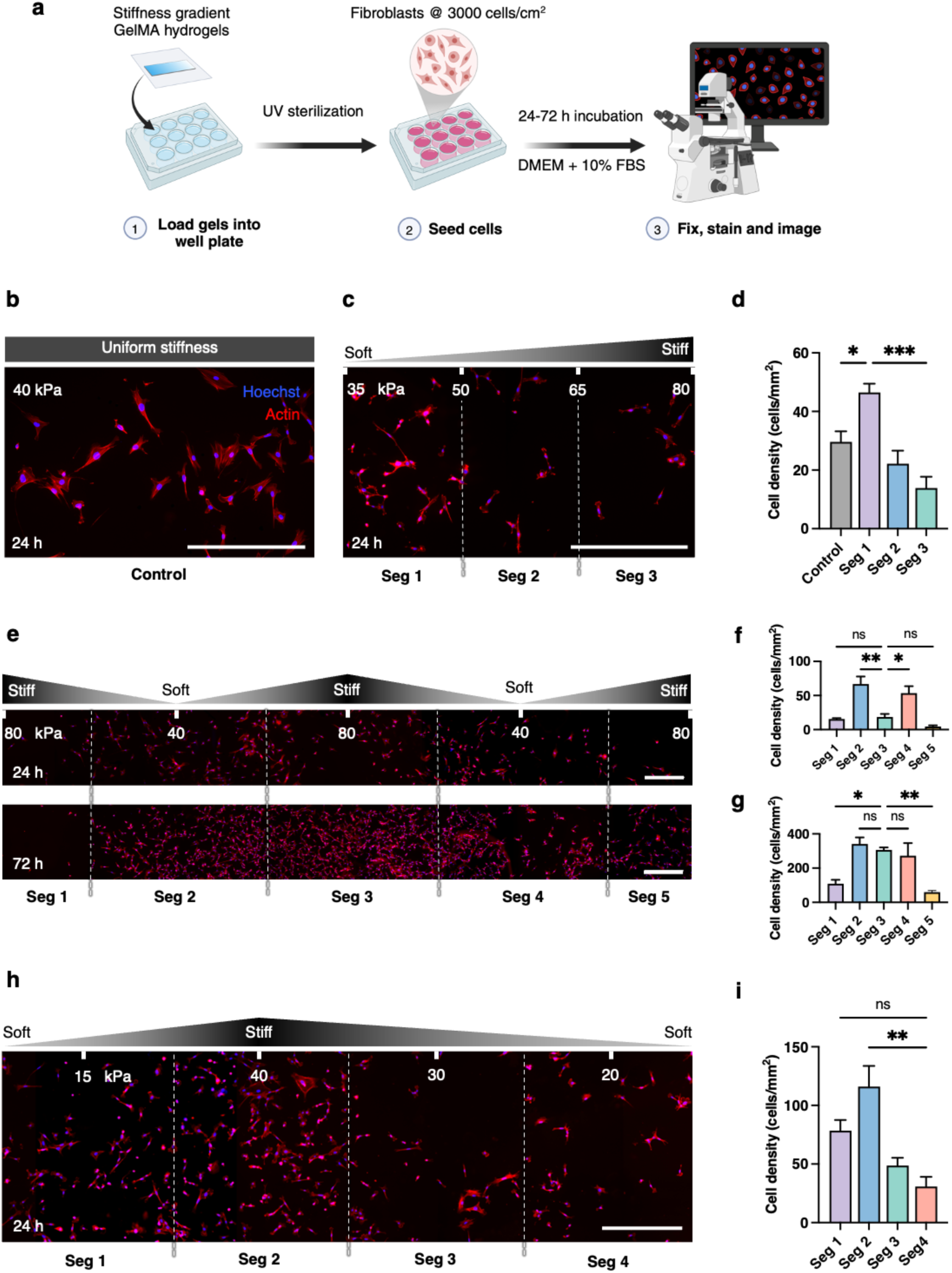
(a) Schematic of experimental process to examine cell behavior of 3T3-L1 fibroblasts on GelMA hydrogels with different stiffness gradient profiles. Representative images of fibroblasts on (b) control sample of uniform stiffness and (c) linear stiffness gradient with a steep slope ≈28 Pa µm^−1^ at 24 h. Quantification of cell distribution on the control and linear gradient gels are shown in (d). (e) Representative images of fibroblasts on symmetrical, periodic stiffness gradient with a slope of linear part ≈16 Pa µm^−1^ at 24 h (top) and 72 h (bottom). (f) Quantification of cell distribution on the periodic gradient gels at 24 h and (g) 72 h. (h) Representative image of fibroblasts on anisotropic stiffness gradient comprising a steep positive slope (≈22 Pa µm^−1^) and a shallow negative slope (≈11 Pa µm^−1^). (i) Quantification of cell distribution on the dual-gradient gels at 24 h. Dashed lines indicate the boundary between segments along the imaged samples for cell analysis. Scale bars: 500 µm. Error bars represent standard error.

A stiffness-dependent cell distribution on the hydrogels was observed after 1 day of culture. As expected, cells on the control sample with uniform stiffness (≈40 kPa) did not exhibit a biased distribution in any direction (**Figure 6b**). On the other hand, the fibroblasts exhibited a graded cell distribution on the linear (**Figure 6c,d**) and periodic gradient gels (**Figure 6e,f**), with a higher population observed toward the softer regions. In contrast, cells accumulated at the stiffer region on the dual-gradient gels (**Figure 6h,i**). The higher local cell densities were interpreted in terms of stiffness-mediated attachment, proliferation, and directional migration. Interestingly, when examining fibroblasts cultured on the periodic gradient gels for 3 days, there appeared to be a biased shift in cell distribution toward the center where the hydrogel stiffness peaks at ≈80 kPa (**Figure 6e,g**). Further analysis revealed that the increased cell density on day 3 relative to day 1 was overall more pronounced in the stiff regions (segments 1, 3 and 5) than the soft regions (segments 2 and 4) (**Figure S4**). Taken together, these results suggest that the initial cell distribution was mainly influenced by a preferential attachment and proliferation around the 30-40 kPa stiffness range, but a gradient slope of >10 Pa µm^−1^ was sufficient to induce durotactic migrate on of fibroblasts, thus resulting in a shift in cell distribution over time.

Overall, the absolute value of stiffness alone does not fully explain cell behavior. Instead, the pattern of stiffness change and the gradient slope can synergistically influence cell-material interactions in profound ways.^[43,44]^ To unravel these complexities, a biomaterials platform such as that developed here can present various complex gradient profiles to provide more information on the cell-material interplay compared to solely testing on linear stiffness gradients, making it a powerful platform to study cell mechanotransduction on stiffness gradients.

## 3. Discussion

The stiffness of the cellular microenvironment strongly correlates with and regulates many biological processes, from cell migration during morphogenesis^[45]^ to matrix remodeling during wound healing^[46]^ and cancer cell invasion.^[47]^ In this context, the development of biomimetic stiffness gradient hydrogels remains an active area of research, driven by the need for systems that more closely resemble the *in vivo* conditions to study mechanosensing and mechanotransduction. Linear stiffness gradients created using polyacrylamide hydrogels have been successfully utilized to uncover fundamental insights into the mechanisms of cell-ECM interplay.^[17,48]^ The univariate elastic mechanics of polyacrylamide gels along with an accessible stiffness range of 0.1-200 kPa were particularly crucial to interrogating the isolated effect of substrate elasticity on cell responses. However, as the field advances in pursuit of a deeper understanding of how cells interpret matrix cues in ever more complex environments, it is anticipated that the applicability of stiffness gradient hydrogels will be improved by the ability to simultaneously modulate other mechanical, structural or biochemical cues within a single platform. Emerging concepts include additional consideration of viscoelastic properties,^[49,50]^ mimicking the anisotropic biophysical gradients within specific tissue niches,^[3]^ and incorporation of fibrous structures to facilitate long-range force transmission,^[51]^ which will serve important functions for future disease modeling and tissue engineering applications.

This work is significant in that it presents a potential solution to this technological challenge by developing a universal platform that can recreate biologically relevant and high-fidelity gradients when coupled with different hydrogel chemistries. Using microfluidics, we demonstrated precise thermal patterning down to micron scale resolution, which could be leveraged to actively manipulate the redistribution of solutes in solution through a process known as thermophoresis.^[52]^ The key accomplishment of this study is demonstrating the feasibility of a modular approach to flexibly achieve user-defined gradient profiles that correlate with a variety of relevant normal and diseased tissue types. Importantly, the reproducibility of stiffness gradients created using this thermophoretic technique was found to depend mostly on the stability of the imposed temperature gradient throughout the specified process time. Provided the polymer solution is completely crosslinked within several minutes after removing the applied temperature gradient, the crosslinking process had minimal impact on the measured variation between samples. Thus, the platform provides a new method to create simple and complex biophysical gradients in soft hydrogels that is decoupled from the crosslinking process. This contrasts with current mainstay techniques based on patterned photo-crosslinking, whereby nonhomogeneous diffusion of radicals and light diffraction can compromise the resulting gradient fidelity in unpredictable ways.^[12]^

Another highlight of the thermophoretic-based gradient hydrogel platform is its compatibility with most of the commonly employed crosslinking mechanisms, as shown with thermally crosslinked GG and photopolymerized GelMA hydrogels in this study. The platform should also be compatible with ionic-crosslinked hydrogels, for example, alginate.^[24]^ This generalizability is attributed to the fact that thermophoresis is a universal physical phenomenon that occurs in any aqueous system when subjected to a non-uniform temperature field.^[53]^ Specifically, the platform leverages thermophoresis effect to establish local variations in concentration within the hydrogel precursor solution. Hydrogel crosslinking is only initiated after the thermophoresis process is complete, meaning that the mechanism of stiffness gradient generation is independent of the crosslinking modality or polymer type. This capability may allow researchers to explore new biomaterial chemistries, enabling the spatial manipulation of hydrogel properties with additional levels of complexity. This part of the study aimed to demonstrate engineered hydrogels possessing a comparable stiffness gradient but varying architecture. It is known that GG exhibits a fibrous architecture whereas GelMA exhibits a honeycomb-like structure.^[22]^ Similarly to how ECM ligands were found to differentially influence cellular responses to substrate stiffness,^[54]^ the matrix architecture might modulate cell mechanotransduction on stiffness gradients. However, these processes have been difficult to investigate due to the lack of reliable strategies that can systematically study and isolate the effects of these two critical ECM properties on cell behavior. Considering the modular approach, the proposed gradient hydrogel platform could facilitate a streamlined fabrication and expand the catalogue of advanced stiffness gradient hydrogel systems for mechanobiology research.

The platform utility was validated by examining fibroblast behaviors on GelMA gradient hydrogels with varying stiffness gradient profiles. The findings highlight how the gradient pattern can be as important in influencing cell behaviors as other characteristics, including absolute stiffness and gradient strength. Although this study employed high system temperatures of average 60 ℃ to accommodate GG processing (critical gelation temperature ≈ 40 ℃),^[22]^ this reflects the needs of the specific material rather than a limitation of the platform. In principle, the thermophoresis-based biofabrication approach can also be implemented at milder conditions using hydrogels that remain liquid at physiological or sub-physiological temperatures, such as GelMA, polyacrylamide, and collagen. To that end, effective thermophoretic gradient generation does not require high absolute temperatures, but rather the presence of a well-defined temperature gradient.^[55]^ This flexibility opens the opportunity for direct fabrication of cell-laden gradient hydrogels, especially when employing smaller temperature gradients. As 3D cell culture systems continue to gain traction for their ability to better mimic the dynamic *in vivo* tissue microenvironment,^[9]^ future studies could leverage the platform introduced here to explore alternative combinations of hydrogel chemistries and thermal conditions that are tailored to the intended cell encapsulation application.

Overall, microengineered stiffness gradient hydrogels holds promising potential for *in vitro* mechanobiology applications. We envision that the proposed modular fabrication strategy can complement the existing biomaterials toolbox and drive the transition of stiffness gradient hydrogels from a niche research tool to a mainstream experimentation platform in mechanobiology research. Of note, stiffness gradient hydrogels have already proven to be valuable for enabling high-throughput screening of cell-material interactions, or used as a reductionist platform to understand the independent effect of a particular material cue on cell responses.^[6,21]^ Patterning of these gradient materials with more complex biophysical landscapes, such as anisotropic stiffness gradients, could further expand their utility in the development of advanced disease models and cell-based hydrogel therapies.^[56,57]^

## 4. Conclusion

This work presented a versatile biofabrication strategy realized by a thermophoretic manipulation approach to creating stiffness gradient hydrogels with a broad stiffness range and gradient strength. The work demonstrated a generalizable strategy to control the microscale mechanical properties in many easily accessible hydrogel systems for cell culture. The capability to pattern simple linear and complex gradients using this modular approach provides a potentially powerful tool to better replicate the spatial mechanical variations within specific tissue niches. The microfabricated platform and the set of characterization techniques demonstrated here can be readily adapted by bioengineers and biologists for use not only to study single-cell mechanobiology, but increasingly to probe ECM-dependent behavior of multicellular structures or cell collectives. Our gradient hydrogel platform promises to achieve designer biomaterials and will contribute to advancing the frontier of research in mechanobiology.

## 5. Experimental Section

### Fabrication and Operation of Thermal Microfluidic Device

Polydimethylsiloxane (PDMS, Sylgard 184, Dow Corning) microfluidic devices were fabricated by replica molding from SU-8 masters based on established soft lithography protocols.^[58]^ The master molds were fabricated by standard photolithography technique on 3-inch silicon wafers (ProSciTech), using high-resolution film photomask (Micro Lithography Services Ltd, UK) and a custom-built UV exposure system.^[59]^ For device assembly, the PDMS replicas were treated using air plasma and bonded to No. 1 coverslips (24 x 50 mm, Trajan Scientific). The device consisted of 9 parallel channels with cross-sectional dimensions of 1 mm (width) × 100 µm (height), whereby adjacent microchannels were separated by a 100 µm-thick PDMS wall running along the length of the channel (**Figure S1**). Depending on the desired thermal pattern, the microchannels were selectively opened to allow water flow for cooling or filled with low melting point alloy (LMPA, Rose’s A alloy, composed of 54% Bi, 28% Pb, 18% Sn) to create micro-Joule heaters for heating. Finally, a sample microchamber was made by cutting out a 1 cm x 0.5 cm area on a 120 µm-thick spacer (2 layers of 60 µm Kapton tape, Multicomp Pro), which was then attached to the underside of the coverslip aligned with the microchannels. For thermophoretic fabrication of gradient hydrogels, the desired temperature gradient across the microfluidic system was maintained by controlling the water flowrate using a syringe pump (DK Infusetek) in tandem with the current imposed across the Joule heater using a power supply unit (Multicomp Pro), as described previously.^[22,24]^

### Temperature Measurement

Temperature distribution across the microfluidic system was characterized using a temperature-sensitive fluorescent dye, Rhodamine B (Sigma-Aldrich).^[29]^ 0.1 mM Rhodamine B solution was drawn into a glass micro-capillary (G346-0101-50, ProSciTech), sealed with epoxy, and attached to the glass coverslip side of the microfluidic device. A thin layer of thermal paste (Chemtronics) was used to hold the micro-capillary in place and facilitate heat conduction. For each thermal patterning condition, the fluorescence intensity imaged during application of temperature gradient was used to generate a heat map based on the established temperature-intensity calibration curve (**Figure S2**).

### Fabrication of Stiffness Gradient Hydrogels

GG precursor solution was prepared by dissolving 1% w/v Gelzan powder (Sigma-Aldrich) in milliQ water at 72 °C and under continuous stirring. Once GG had completely dissolved, 5 mM calcium chloride (CaCl_2_, Sigma-Aldrich) was added as crosslinkers. GelMA precursor solution were prepared by dissolving 5% w/v or 10% w/v freeze-dried GelMA in PBS at 50 °C and containing 0.1% w/v Irgacure 2959 (Sigma-Aldrich). The prepared solution was wrapped in aluminum foil, stored at 4 °C overnight, and used within 1 week. To synthesize GelMA, gelatin was reacted with methacrylate anhydride (MA) as previously described.^[26,31]^ Briefly, 7 g gelatin powder (Type A, ∼175 Bloom, Sigma-Aldrich) was dissolved in 70 mL PBS at 50 °C in a water bath. Then, 5.6 mL MA was added at a rate of 0.5 mL min^−1^ under vigorous stirring. The mixture was left to react for 2 h and the resulting solution was dialyzed against milliQ water for 7 days at 40 °C and finally freeze-dried to obtain a white solid product. The GelMA used in this study had a degree of functionalization of 85%, as determined by ^1^H-NMR analysis.

To fabricate stiffness gradient hydrogels, the thermal microfluidic device was first preheated at the desired water flowrate and controlled current for 5 min to achieve a stabilized temperature gradient. The average system temperature was also maintained above 60 °C using a Peltier module (RS Components) attached to the PDMS layer, which was critical to reduce viscosity of the precursor solutions and prevent premature gelation. 20 µL precursor solution was pipetted into the microchamber and immediately covered with 0.1% poly(ethyleneimine) (PEI)-treated coverslip for GG solution or 3-(trimethoxysilyl) propyl methacrylate (TMSPMA)-treated coverslip for GelMA solution. Then, the applied temperature gradient was maintained for 30-40 minutes to allow thermophoretic generation of a concentration gradient within the micro-confined volume of precursor solution. The resulting concentration gradient was “frozen” by switching the polarity of the Peltier module to enable rapid cooling of the whole system. For GG, the system was kept at approximately 15 °C for 1.5 min to ensure complete gelation. For GelMA, the system was further exposed to UV (365 nm, 0.22 W cm^−2^) for 120 s to form covalently crosslinked hydrogels. Finally, the hydrogel attached to the functionalized coverslip was gently peeled off and stored immersed in PBS at 4 ℃ until use for mechanical characterization or cell study.

### Coverslip Functionalization

PEI-coated coverslips enable attachment of GG hydrogels via electrostatic interaction. Glass coverslips (12 mm x 12 mm, No.1, Marienfield) were cleaned of organics and surface-activated by plasma treatment for 90 s. The coverslips were immediately immersed in 0.1% w/v aqueous PEI (03880, Sigma-Aldrich) solution and allowed to coat overnight at room temperature. After that, the coverslips were rinsed under a stream of milliQ water and air dried. The PEI-coated coverslips were stored at room temperature in a dry environment, used within 1 month.

Methacrylated coverslips enable attachment of GelMA hydrogels via covalent bonding when exposed to UV. Glass coverslips (12×12 mm, No.1, Marienfield) were plasma treated for 90 s and then immersed in a 200:6:1 solution of absolute ethanol, acetic acid, and TMSPMA (440159, Sigma-Aldrich) for 5 min. Then, the coverslips were washed in absolute ethanol for 3 min and air dried. The TMSPMA functionalized coverslips were stored at room temperature in the dark, used within 2 weeks.

### Stiffness Characterization

Stiffness (Young’s Modulus, kPa) across the fabricated gradient hydrogels were characterized via nanoindentation using an MFP-3D atomic force microscopy (Asylum Research), following an established protocol^[60]^ with minor adaptations. Samples were indented using 10 µm diameter, borosilicate glass colloidal probe, with 30 kHz resonance frequency and 0.1 N m^−1^ nominal spring constant (CP-qp-CONT-BSG-B-5, NanoAndMore). The cantilever spring constant was more precisely determined using the thermal calibration method at the start of each experiment session. Samples were immersed in a drop of PBS, indented at an approach velocity of 2 µm s^−1^ until a 15 nN trigger force was registered, then retracted at 10 µm s^−1^ (**Figure S5**). Measurements were taken at 200 µm intervals for linear and sawtooth gradient hydrogels, and at 500 µm intervals for extended linear and periodic gradient hydrogels. Each measurement point represents the average Young’s modulus of a force map of 16 force curves generated over a 20 µm x 20 µm area. The force curves were analyzed directly in the Asylum MFP-3D software using the Hertz model.

### Cell Culture

Mouse fibroblasts (3T3-L1) were cultured in high-glucose DMEM (Gibco) supplemented with 10% v/v fetal bovine serum (Gibco), and 100 U mL^−1^ Penicillin/Streptomycin (Gibco) at 37 °C and 5% CO_2_. Cells were split using TrypLE Express (Gibco) and used up to passage 20 for experiments. Hydrogels were sterilized with UV-C light in PBS for 30 min prior to cell seeding. For all experiments, 3T3-L1 cells were seeded on the gels at 3000 cells cm^−2^. The cells were allowed to attach on the gels for 1 h before additional media was added, followed by incubation at 37 °C and 5% CO_2_ for 24 hr. For periodic gradient hydrogels, an additional study was performed where the cells were cultured under the same conditions for 72 h. As a control experiment, 3T3-L1 cells were cultured onto uniform stiffness GelMA hydrogels prepared using 10% w/v GelMA and 0.1% w/v Irgacure 2959.

### Immunofluorescence Staining and Imaging

Cells were fixed using 4% paraformaldehyde for 20 min at room temperature and washed (3 x 5 min) with PBS-T (Phosphate Buffer Saline with 0.5% Triton X-100) to permeabilize the cells. Cells were stained with rhodamine phalloidin (94072, Sigma-Aldrich) and Hoechst 33342 (14533, Sigma-Aldrich). Samples were imaged on an inverted fluorescence microscope (IX73, Olympus) using a 10x objective. All images were acquired using a highly sensitive fluorescent camera (Prime BSI Express, Teledyne Photometrics) interfaced with Micro-Manager open-source microscopy software. For each gradient hydrogel, an image mosaic was generated by stitching multiple images acquired along the length of the gradient, using DAPI as the reference channel. Resulting stitched images were then cropped into multiple segments representing regions of different stiffness along the hydrogel, then analyzed using FIJI software. For each segment, cell nuclei were detected as objects through manual thresholding and counted using the ‘Analyze Particle’ tool. The cell density was then calculated by dividing the total number of cells by the hydrogel segment area.

### Statistical Analysis

Data were presented as mean ± standard error of the mean (SEM). All experiments were performed in triplicates. Statistical tests were performed on GraphPad Prism (v10) using a one-way ANOVA with Tukey post-hoc test. Statistical significance between experimental groups of interest were denoted as * (p < 0.05), ** (p < 0.01), *** (p < 0.005), or *ns* (non-significant).

## Acknowledgements

S.W.C. acknowledges the University of Sydney Faculty of Engineering Research Scholarship for financial support. The authors thank A/Prof. Sara Baratchi for insightful advice and discussions on cell culture experiments; Dr. Chris Vega-Sanchez for helpful suggestions on spacer design; Jiayan Shao and Bingyan Liu for advice on GelMA synthesis. J.C. and G.M. were supported by the Sydney Nano Taste of Research Award.

## Author Contributions

S.W.C designed the research, conducted the experiments, analyzed the data, and wrote the original manuscript. J.S. assisted with the cell culture experiments. G.M. performed preliminary experiments to optimize the microfluidic system design. J.C. contributed to the fabrication of stiffness gradient hydrogel samples and cell data analysis. A.K. and I.J.G contributed to the methodology development for AFM experiments. M.B., Y.S., and C.N. provided supervision, and acquired resources and funding. D.V. designed the research, provided supervision, and acquired resources and funding. All authors have reviewed and approved the final version of the manuscript.

